# Genome-wide association analysis of excessive daytime sleepiness identifies 42 loci that suggest phenotypic subgroups

**DOI:** 10.1101/454561

**Authors:** Heming Wang, Jacqueline M Lane, Samuel E Jones, Hassan S Dashti, Hanna Ollila, Andrew R Wood, Vincent T. van Hees, Ben Brumpton, Bendik S Winsvold, Katri Kantojärvi, Brian E Cade, Tamar Sofer, Yanwei Song, Krunal Patel, Simon G Anderson, David A Bechtold, Jack Bowden, Richard Emsley, Simon D Kyle, Max A Little, Andrew S Loudon, Frank AJL Scheer, Shaun M Purcell, Rebecca C Richmond, Kai Spiegelhalder, Jessica Tyrrell, Xiaofeng Zhu, Kati Kristiansson, Sonja Sulkava, Tiina Paunio, Kristian Hveem, Jonas B Nielsen, Cristen J Willer, John-Anker Zwart, Linn B Strand, Timothy M Frayling, David Ray, Deborah A Lawlor, Martin K Rutter, Michael N Weedon, Susan Redline, Richa Saxena

**Author notes:** These authors contributed equally (Deborah A Lawlor, Martin K Rutter, Michael N Weedon, Susan Redline, Richa Saxena). Corresponding author* R. Saxena, Center for Genomic Medicine, Massachusetts General Hospital, 185 Cambridge Street, CPZN 5.806, Boston, MA, 02114, USA.

## Abstract

Excessive daytime sleepiness (EDS) affects 10-20% of the population and is associated with substantial functional deficits. We identified 42 loci for self-reported EDS in GWAS of 452,071 individuals from the UK Biobank, with enrichment for genes expressed in brain tissues and in neuronal transmission pathways. We confirmed the aggregate effect of a genetic risk score of 42 SNPs on EDS in independent Scandinavian cohorts and on other sleep disorders (restless leg syndrome, insomnia) and sleep traits (duration, chronotype, accelerometer-derived sleep efficiency and daytime naps or inactivity). Strong genetic correlations were also seen with obesity, coronary heart disease, psychiatric diseases, cognitive traits and reproductive ageing. EDS variants clustered into two predominant composite phenotypes - sleep propensity and sleep fragmentation - with the former showing stronger evidence for enriched expression in central nervous system tissues, suggesting two unique mechanistic pathways. Mendelian randomization analysis indicated that higher BMI is causally associated with EDS risk, but EDS does not appear to causally influence BMI.

---

EDS is a chief symptom of chronic insufficient sleep ^1^ as well as of several primary sleep disorders, such as sleep apnea, narcolepsy, and circadian rhythm disorders ^2–5^. Several disease processes and medications also associate with prevalent and incident EDS ^6–9^. EDS is estimated to contribute to risk for motor vehicle crashes, work-related accidents and loss of productivity, highlighting its public health importance ^10,11^. The clinical impact of EDS extends to a negative impact on cognition, behavior, and quality of life ^12,13^. Therefore, sleep interventions often identify reduction in EDS as a chief goal ^14,15^. EDS is also associated with an increased risk for cardio-metabolic disorders, psychiatric problems and mortality ^9,16,17^ through pathways that may be causal, bi-directional, or reflect pleiotropic effects.

While EDS occurs in a variety of settings associated with insufficient sleep, there is large inter-individual variability in levels of EDS that is not fully explained by sleep duration, sleep quality or chronic disease. Experimental studies have shown that there is also individual vulnerability to EDS following sleep restriction ^18,19^. The heritability of EDS is estimated to be between 0.37 to 0.48 in twin studies, 0.17 in family studies, and between 0.084 to 0.17 in GWAS ^20,21^, suggesting that genetic factors contribute to variation in sleepiness. Despite multiple candidate gene studies ^22–24^ and GWAS ^20,25,26^, including one from the first genetic release of the UK Biobank ^21^, few significant genetic variants have been reported, likely reflecting the heterogeneous and multifactorial etiology of the phenotype and low statistical power. Here, we extend our EDS GWAS to the full UK Biobank dataset ^27^, enabling identification of multiple genetic variants and molecular pathways that may contribute to EDS.

In the UK Biobank, ^27^ 452,071 participants of European genetic ancestry self-reported frequency of EDS using the question: “How likely are you to doze off or fall asleep during the daytime when you don’t mean to? (e.g.: when working, reading or driving)”, with the answer categories “never” (N=347,285), “sometimes” (N=92,794), “often” (N=11,963), or “all of the time” (N=29). The severity of EDS increased with older age, female sex, higher BMI, various behavioral, social and environmental factors, and chronic diseases (Supplementary Table 1). EDS was positively correlated with self-reported insomnia symptoms, morning chronotype, ICD10 or physician diagnosed sleep apnea and self-reported shorter and longer sleep duration, consistent with earlier reports or known clinical correlates ^21^ (Supplementary Table 1a and 2). EDS was also correlated with shorter sleep duration, lower sleep efficiency (indicating more time awake during the sleep period) and longer daytime inactivity duration estimated using a 7-day accelerometry in a subset (N=85,388) of UK Biobank participants (Methods; Supplementary Table 1b) ^20^.

## GWAS and replication

We performed a GWAS of EDS treating the four categories as a continuous variable using a linear mixed regression model^28^ adjusted for age, sex, genotyping array, ten principal components (PCs) of ancestry and genetic relatedness matrix, and identified 37 genome-wide significant loci (p<5×10^−8^) (Fig. 1, Supplementary Fig.1 and Supplementary Table 3). Previously identified loci for EDS in the first release of the UK Biobank (N=111,975) ^21^, including loci at or near *PATJ, HCRTR2*, and *CPEB1*, were confirmed in this study. The most significant association was observed at *KSR2*, a gene regulating multiple signaling pathways, including the ERK/MEK signaling pathway, affecting energy balance, cellular fatty acid and glucose oxidation that is implicated in obesity, insulin resistance and heart rate during sleep in previous studies in humans and mice ^29,30^. Additional novel loci were identified within or near genes with known actions on sleep-wake control regulation or that are associated with sleep disorders (e.g. *PLCL1^31^, GABRA2 ^32^, BTBD9 ^33^, HTR7 ^34^, RAI1^35^)*, metabolic traits (e.g. *GCKR ^36^, SLC39A8 ^37^)* and psychiatric traits (e.g. *AGAP1^38^, CACNA1C^39^)*. No association was seen with SNPs highlighted in smaller independent GWAS of EDS, hypersomnia or narcolepsy ^20,25,26,40–45^ (Supplementary Table 4).

**Figure 1.**
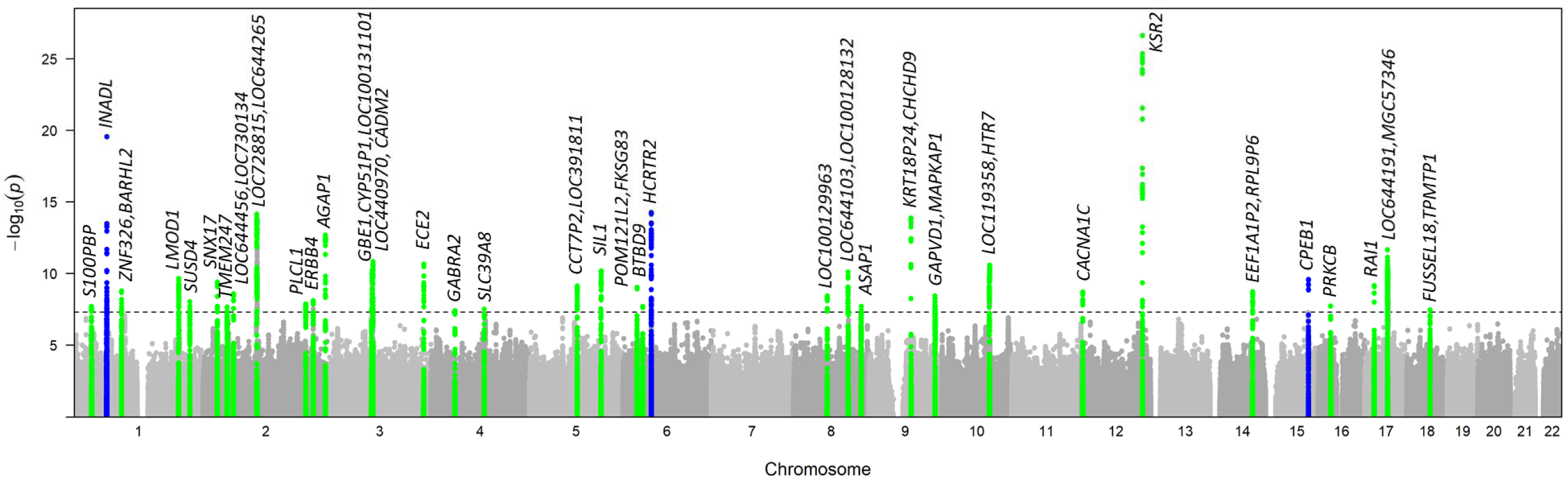
Genome-wide association analysis of EDS (modelled as a 4-level continuous variable) identified 37 significant loci (P<5×10^−8^), including previously associated signals near *PATJ, HCRTR2, and CPEB1* (blue). Genes near novel significant loci are highlighted in green.

Previous longitudinal research indicated obesity and weight gain were associated with EDS incidence ^8^; therefore, we performed an additional GWAS adjusting for BMI to identify loci that may operate in obesity-independent pathways. This analysis identified 5 additional loci (Supplementary Fig. 2 and Supplementary Table 3). Effect estimates at the 37 loci identified in the primary model were largely unchanged.

Sensitivity analyses adjusting for potential confounders (including depression, socio-economic status, alcohol intake frequency, smoking status, caffeine intake, employment status, marital status, and psychiatric problems) did not substantially alter effect estimates of the 42 identified signals (Supplementary Table 5). Analyses stratified by obesity and sleep duration revealed consistent effect directions (heterogeneity P>0.05) but lower statistical evidence of association, likely reflecting the smaller subgroup sample sizes (Supplementary Table 6). Secondary GWAS (N=255,426), excluding shift workers and individuals with chronic health or psychiatric illnesses, additionally identified significant variants in *SEMA3D* and revealed marginal significant interactions with health status at *PATJ, ZENF326/BARHL2, ECE2, ASAP1*, and *CYP1A1/CYP1A2* (p<0.05; Supplementary Fig. 3 and Supplementary Table 7). Conditional analyses at each of the 42 loci identified no secondary signals. Sex-stratified analyses did not reveal significant gene by sex interactions (Supplementary Fig. 4 and Supplementary Table 8).

Replication was attempted using self-reported (using different questions to ours and each other) EDS indices available in European whites from HUNT ^46^ (N=29,906), FINRISK ^47^ (N=20,344), and Health 2000 studies ^48^ (N=4,546) (Methods; Supplementary Table 9). Only eight individual signals, including *KSR2*, were marginally significant (p<0.05) in individual cohorts and/or meta-analysis of these, likely due to the lower power, inconsistent questionnaires across different cohorts and the multi-factorial etiology of EDS (Supplementary Table 10). However, a genetic risk score (GRS) of all 42 EDS loci weighted by the effect estimates from our primary EDS GWAS was associated with increased EDS in a meta-analysis of HUNT, FINRISK and Health 2000 (Fisher’s p=0.001; Supplementary Table 10).

## Heterogeneous effects of EDS loci on other sleep traits

The 42 EDS-associated genetic variants likely influence EDS through different mechanisms. Therefore, to dissect heterogeneity, we tested for association of the combined GRS and individual SNPs with subjective and objective measures of sleep patterns and disorders (Table 1 and Supplementary Table 11).

**Table 1.**
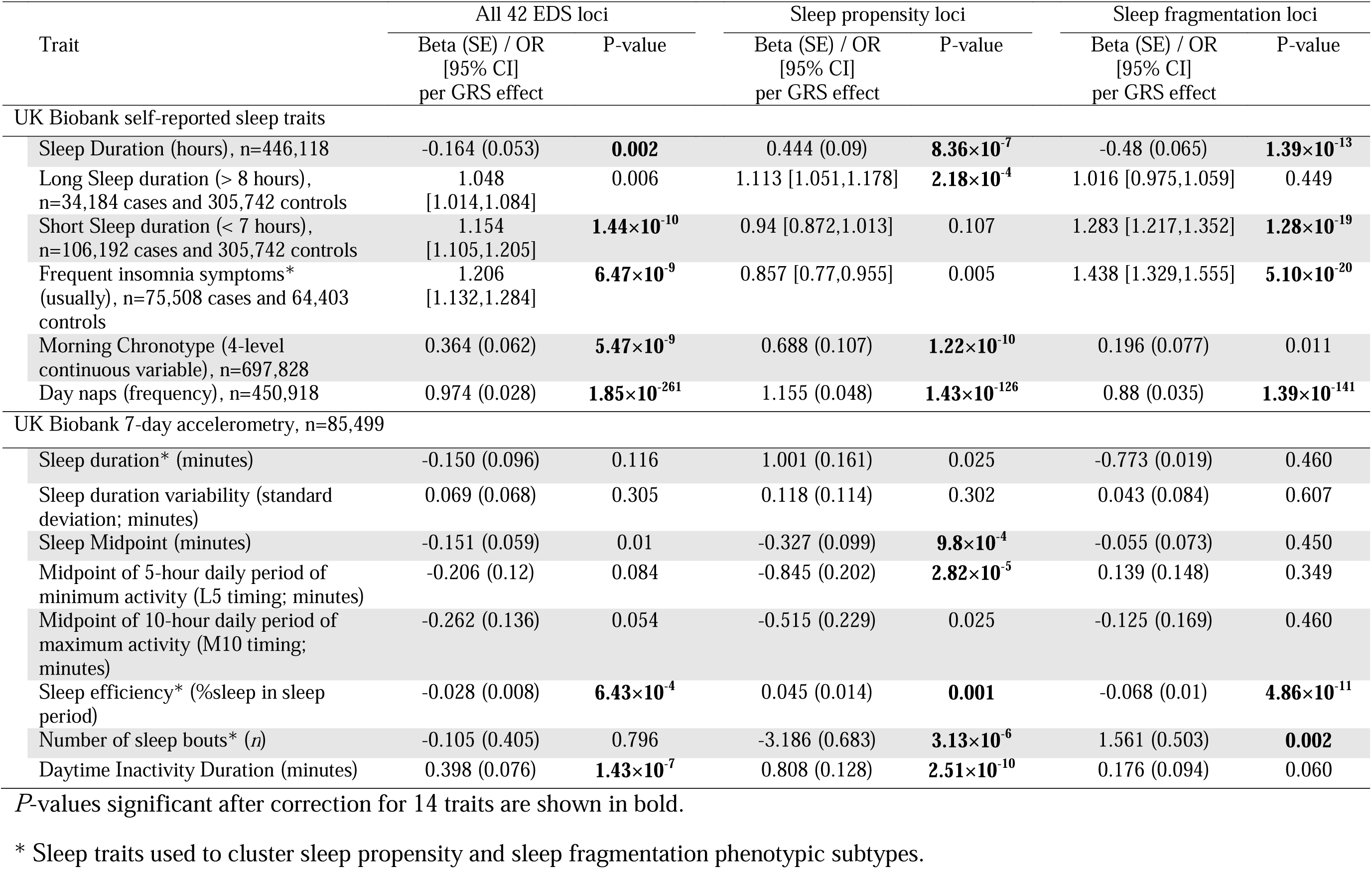
Association of EDS weighted genetic risk score (GRS) of all 42 EDS loci, sleep propensity loci, and sleep fragmentation loci with a) multiple self-reported sleep traits and b) 7-day accelerometry-derived sleep, circadian, and activity traits in the UK Biobank.

A GRS of all 42 variants was associated with self-reported shorter sleep duration, morning chronotype, increased insomnia risk, increased frequency of daytime napping, and with accelerometry-assessed lower sleep efficiency and increased duration of daytime inactivity (Table 1). The EDS GRS was not associated with 7-day accelerometry-derived continuous sleep duration, which may reflect differences in self-reported habitual sleep vs. accelerometry-estimated sleep, different time points in data collection (accelerometry data were collected 5-7 years after questionnaires), lower power given smaller sample size in accelerometry data, or recall or other biases in the self-report such as from reporting in 1 hour increments.

Individual EDS increasing alleles at *PATJ* and *PLCL1* were also associated with morning chronotype (as previously reported^31,49^); at metabolism regulatory genes *KSR2*, *LOC644191*/*CRHR1* and *SLC39A8* with self-reported sleep duration (*KSR2* with increased sleep duration, *LOC644191/CRHR1* with long sleep and *SLC39A8* with short sleep); *LMOD1* and *LOC644456/LOC730134* with both insomnia and short sleep duration; and at the orexin/hypocretin receptor *HCRTR2* (known to play a central role in sleep wake control and narcolepsy ^50^) with both morning chronotype and short sleep duration, suggesting common genetic factors. Consistently, adjusting for sleep disturbance traits (ICD10 code defined sleep apnea or narcolepsy, or self-reported sleep duration hours, frequent insomnia symptoms or chronotype) together attenuated effect estimates for several loci, suggesting that these genetic variants influence EDS through altered sleep patterns; however, adjustment for any trait alone only minimally altered effect estimates at select individual loci (Supplementary Table 5).

Using 7-day accelerometry-derived data available in a subset of the UK Biobank (N=85,388), we observed associations of several EDS alleles with reduced sleep efficiency (e.g., *SNX17*) whereas others were associated with increased sleep efficiency (e.g., *PLCL1*), suggesting that genetic mechanisms may lead to EDS through effects on increased sleep fragmentation (i.e., low sleep efficiency) or increased sleep propensity (i.e., high sleep efficiency), respectively. Therefore, we performed clustering analysis on EDS risk alleles at 42 loci according to their association effect sizes (z-scores) with objective estimates of sleep efficiency, sleep duration, and number of sleep bouts, and self-reported frequent insomnia symptoms. We interpreted EDS alleles showing patterns of association with higher sleep efficiency, longer sleep duration, fewer discrete sleep bouts and fewer insomnia symptoms as reflective of greater sleep propensity; whereas EDS alleles associated with these sleep traits in a largely inverse manner were interpreted as reflective of disturbed sleep or a sleep fragmentation phenotype (Fig. 2). GRS of EDS loci stratified by the two clusters support our interpretation, with sleep propensity loci showing robust associations with early circadian traits, (e.g. morning chronotype P=1.22×10^−10^; Table 1). Future statistically robust clustering analysis approaches including other sleep and related traits will be needed to validate and distinguish the phenotypic subtypes of EDS ^51^

**Figure 2.**
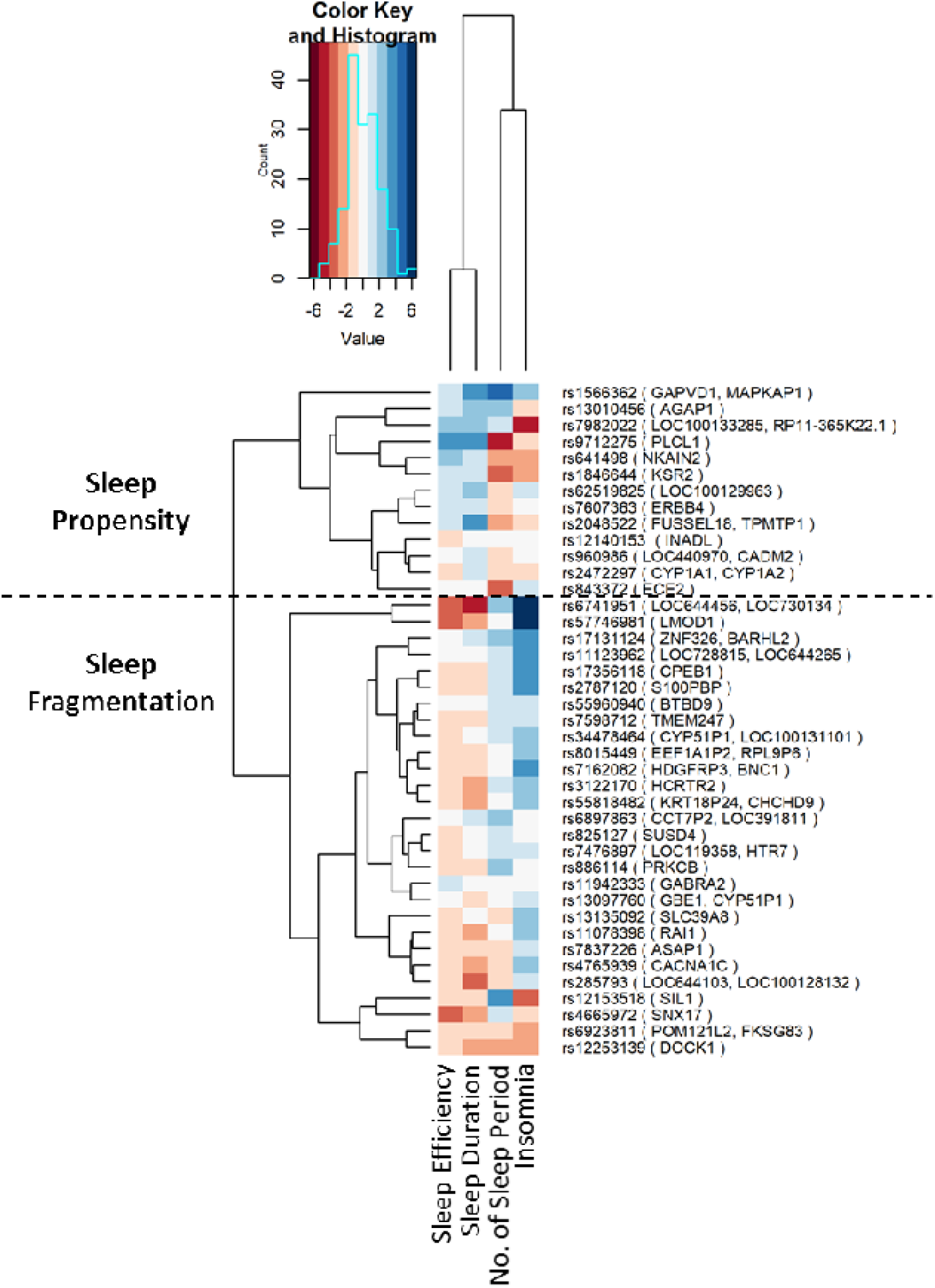
EDS risk alleles associated predominantly with sleep propensity or sleep fragmentation phenotypes. Each cell shows effect sizes (z-scores) of associations between sleep traits (accelerometry-derived sleep efficiency, sleep duration, number of sleep bouts, and self reported insomnia symptoms) and EDS risk alleles (positively associated with EDS). Sleep propensity alleles were defined as more likely associated with higher sleep efficiency, longer sleep duration, fewer sleep bouts and fewer insomnia symptoms. Sleep fragmentation alleles were defined as more likely associated with lower sleep efficiency, shorter sleep duration, more sleep bouts and more insomnia symptoms.

**Figure 3.**
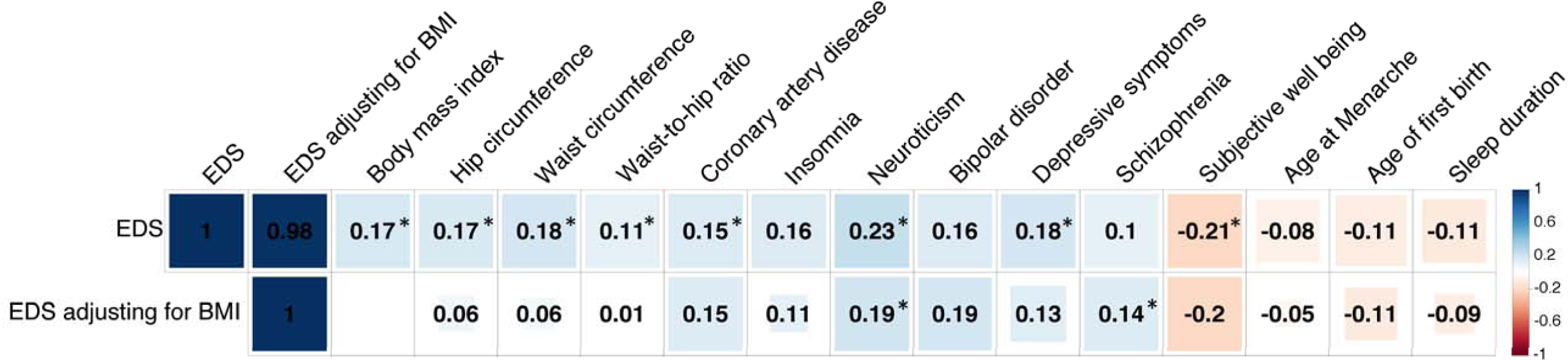
Top significant genetic correlations (r_g_) between EDS and published summary statistics of independent traits using genome-wide summary statistics using LD score regression (LDSC). Blue color indicates positive genetic correlation and red color indicates negative genetic correlation. Larger colored squares correspond to more significant P values, and asterisks indicate significant (P <2.2×10^−4^) genetic correlations after adjusting for multiple comparisons of 224 available traits. All genetic correlations in this report can be found in tabular form in Supplementary Table 20.

**Figure 4.**
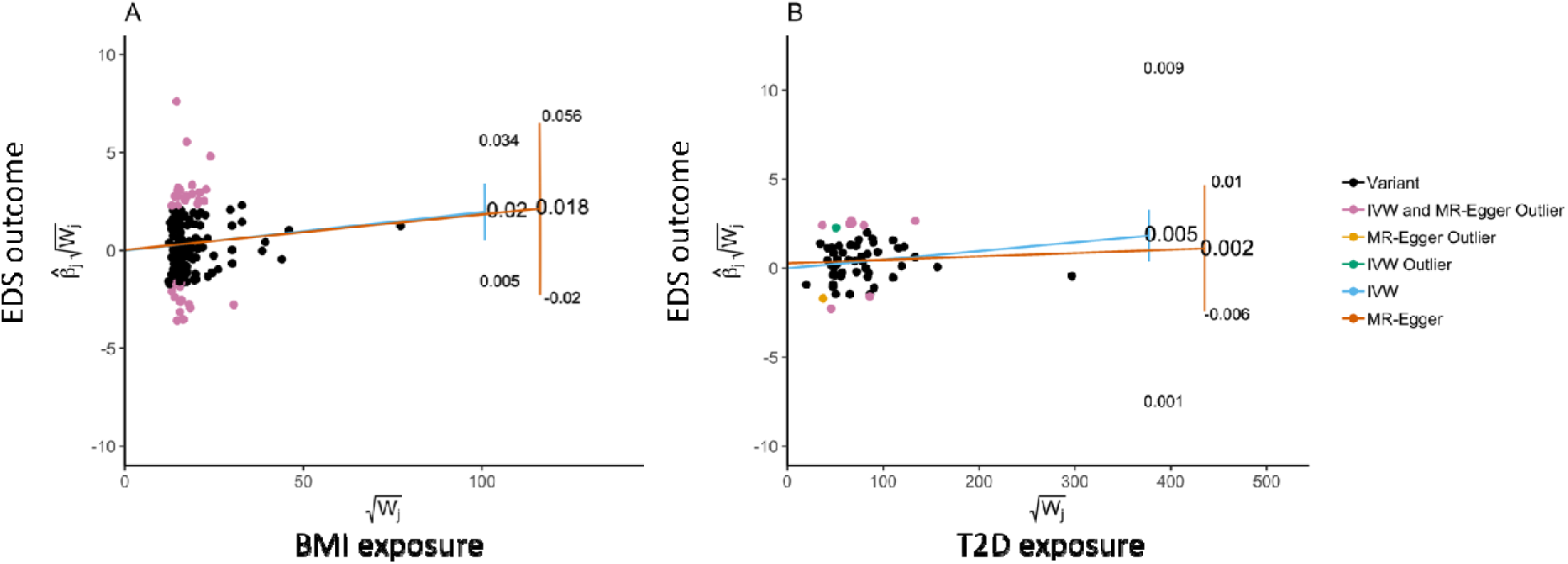
Radial plot of two-sample Mendelian randomization between (A) BMI and (B) Type 2 diabetes with EDS outcome using IVW and MR-Egger tests. The x-axis is the inverse standard error (square root weights in the IVW analysis) for each SNP. The y-axis scale represents the ratio estimate for the causal effect of an exposure on outcome for each SNP () multiplied by the same square root weight.

## Functional effects of loci

The 42 loci lie in genomic regions encompassing 164 genes (Supplementary Table 12), and 3 associations are in strong linkage disequilibrium with known GWAS associations for other traits, including blood cell count, HDL cholesterol and caffeine metabolism ^52–55^. Genes at multiple loci have been implicated in Mendelian syndromes or in experimental studies in mouse or fly models. Eighteen loci harbor one or more genes with potential drug targets ^56,57^.

We performed fine mapping analyses for potential causal variants using PICS ^58^ and identified 33 variants within 25 EDS loci with a causal probability larger than 0.2 (Supplementary Table 13). The majority of likely causal variants were intronic (65%) followed by non-coding transcript variants (8%) and nonsense mediated decay (NMD) transcript variants (7%) (Supplementary Fig. 5). Functional variants included a missense variant rs12140153 within *PATJ*, a synonymous variant rs11078398 within *RAI1*, regulatory variants rs10800796 in the promoter region of *LMOD1*, and rs239323 in a CTCF binding site in the gene *POM121L2*. Using the Oxford Brian Imaging Genetics (BIG) server ^59^, we further observed the pleiotropic locus at rs13135092 (*SLC39A8 ^60^*) to be significantly associated with bilateral putamen and striatum volume in the UK Biobank (p<2.8×10^−7^; N=9,707; Supplementary Fig. 6). This could be of particular interest given the importance of these central brain centers in influencing motor and emotional behaviors, and emerging data implicating these centers in the integration of behavioral inputs that modulate arousal and sleep-wake states ^61,62^.

## Gene-based, pathway and tissue enrichment analyses

Gene-based analyses using PASCAL ^63^ identified 94 genes associated with EDS (p<2.29×10^−6^) (Supplementary Table 14), of which 63 overlapped with genes under significant association peaks shown in Supplementary Table 13. Tissue enrichment analysis across 53 tissues in the GTEx database^64^ using MAGMA^65^ identified multiple brain tissues including the frontal cortex, cerebellum, anterior cingulate cortex, nucleus accumbens, caudate nucleus, putamen, hypothalamus, amygdala, and hippocampus (P<10^−3^; Supplementary Table 15 and Supplementary Fig. 7A), i.e., including brain regions implicated in sleep-wake and arousal disorders ^66^ as well as centers responsive to sleep deprivation. Pathway and ontology analyses using PASCAL identified enrichment in neuronal synaptic transmission pathways and EnrichR ^67,68^ identified pathways involved with the central nervous system, neurotransmitters, and metabolic processes (e.g. insulin receptor signaling pathway) (Supplementary Table 16 and 17). Genes at loci showing clustering with sleep propensity phenotypes (n=67) showed enriched expression in brain tissues including cortex, hippocampus, cerebellum, and amygdala (P<10^−3^; Supplementary Fig. 7B). In contrast, no tissues were enriched in expression of genes that showed clustering with sleep fragmentation phenotypes (n=97) (Supplementary Fig. 7C), suggesting that the mechanisms associated with sleep fragmentation may be more complex, reflective of multifactorial influences. Pathway and ontology analyses results for clustered genes using FUMA also reveal different patterns (Supplementary Fig. 8 and 9).

The heritability of EDS explained by genome-wide SNPs was estimated at 6.9% (SE=1%). Partitioning heritability across tissue types and functional annotation classes indicated enrichment of heritability in central nervous system and adrenal/pancreas tissue lineage tissues, and in regions conserved in mammals, introns, and H3K4me1 - potentially active and primed enhancers ^69^ (P<8.3×10^−4^) (Supplementary Table 18).

## Genetic overlap among EDS, sleep disorders and other disease traits

Consistent with EDS being a symptom of several sleep disorders, GRSs of genome-wide significant SNPs for restless leg syndrome ^70^ (P=0.0002), insomnia ^71^ (P=4×10^−7^), and coffee consumption ^54^ (P=1.87×10^−12^) (often used as a sleepiness “counter-measure”) were significantly associated with EDS phenotype (Table 2). Although EDS is a key symptom of narcolepsy, the GRS of narcolepsy ^45^ was not associated with EDS phenotype (P=0.126), suggesting narcolepsy loci did not explain EDS variation in this sample. We could not examine the genetic overlap of sleep apnea loci and EDS because few significant loci for sleep apnea have been reported in the literature ^35^ and there was limited sleep apnea information in this cohort.

**Table 2.** Association between weighted genetic risk scores (GRS) of significant SNPs (P<5×10^−8^) for other sleep behavioral traits and sleep disorders with phenotypic EDS in UK Biobank.

| Trait | N | nSNP | Beta (SE)<br>per GRS effect | P-value |
| --- | --- | --- | --- | --- |
| Frequent Insomnia Symptoms <sup>71</sup> | 237,620 | 57 | 0.0007 (0.0001) | **$4.00 \times 10^{-7}$** |
| Sleep Duration (hours) <sup>78</sup> | 446,118 | 78 | -0.0002 (0.0001) | 0.193 |
| Short Sleep <sup>78</sup> | 106,192 | 27 | 0.0009 (0.0002) | **$1.25 \times 10^{-4}$** |
| Long Sleep <sup>78</sup> | 34,184 | 8 | 0.0009 (0.0005) | 0.068 |
| Day Naps | 450,918 | 125 | 0.4078 (0.011) | **$6.61 \times 10^{-281}$** |
| Morning Chronotype <sup>79</sup> | 697,828 | 348 | 0.00004 (0.0001) | 0.524 |
| Restless Leg syndrome <sup>70</sup> | 110,851 | 20 | 0.0009 (0.0002) | **$2.21 \times 10^{-4}$** |
| Narcolepsy <sup>45</sup> | 25,857 | 8 | 0.0007 (0.0004) | 0.126 |
| Coffee Consumption (cups) <sup>54</sup> | 91,462 | 8 | 0.033 (0.005) | **$1.87 \times 10^{-12}$** |

*P*-values significant after correction for 9 traits are shown in bold.

To investigate the genetic correlation between EDS and other common disorders, we tested the proportion of genetic variation of EDS shared with 233 other traits with published GWAS summary statistics in LDSC^72^. After adjusting for multiple comparisons, significant positive genetic correlations were observed for EDS with obesity traits, coronary heart disease, and psychiatric traits (P<0.0001) (Supplementary Table 19). The genetic correlations of EDS with coronary artery disease and psychiatric traits persisted after adjusting for BMI (Figure 2), perhaps partially reflecting shared neurologic or neuroendocrine factors, such as those that underlay insomnia and short sleep with cardiac and psychiatric traits ^73,74^. Consistently suggestive negative genetic correlations for EDS with subjective well-being and reproductive traits (age at menarche and age at first birth) were also observed (P<0.005).

To evaluate the causal relationship between sleep disorders or other disease traits and EDS, we performed two-sample summary-level Mendelian Randomization (MR) analyses using independent genetic variants from published summary statistics from GWAS of BMI, type 2 diabetes, coronary heart disease, neuroticism, bipolar disorder, depression, schizophrenia, age of menarche, restless leg syndrome, narcolepsy, insomnia, sleep duration, and chronotype as exposures and EDS as outcome ^75^. Using the inverse variance weighted (IVW) approach, we identified a putative causal association of higher BMI with increased EDS (β=0.018; 95% CI [0.008, 0.028]; P-value=0.0004; Supplementary Table 20), which was significant after accounting for multiple comparisons (P-value<0.003). However, there was evidence of variant heterogeneity potentially due to horizontal pleiotropy (Cochran’s Q=677.17; P=1.09×10^−37^). Therefore, we performed sensitivity analysis using the Radial MR-Egger approach (Methods)^76^ to control for bias due to pleiotropy, and observed an effect that was consistent with our main IVW analyses but less precisely estimated (wider confidence intervals) because this method is statistically relatively inefficient (β=0.025; 95% CI [−0.005,0.055]; P-value=0.103; Supplementary Table 21). These results support findings from prospective epidemiological studies ^8,77^. An additional suggestive putative causal association of type 2 diabetes with increased EDS was also observed (IVW β=0.005; 95% CI [0.001, 0.009]; P-value=0.014; Supplementary 20) with evidence of heterogeneity (Cochran’s Q=88.38; P=0.005; Supplementary 21), but broadly consistent results when using Radial MR-Egger again showed a consistent effect direction (β=0.002; 95% CI [−0.006,0.01]; P-value=0.637). Reverse MR did not identify any strong evidence for EDS having a causal effect on any of the outcomes we examined (Supplementary Table 22).

In summary, an extensive series of analysis from a large GWAS for EDS has begun to unravel the heterogeneous genetic architecture of EDS and has identified genetic correlations and causal relationships with other diseases and lifestyle traits. Multiple genetic loci were identified, including genes expressed in brain areas implicated in sleep-wake control and genes influencing metabolism. Shared genetic factors were identified for EDS and other sleep disorders and health outcomes. EDS variants clustered with two predominant EDS phenotypes-sleep propensity and sleep fragmentation-with the former showing stronger evidence for enrichment in central nervous system tissues, suggesting two unique mechanistic pathways. Moreover, Mendelian randomization analysis indicated that higher BMI is causally associated with increased EDS.

## Acknowledgements and Funding

This research has been conducted using the UK Biobank Resource under applications 6818 and 9072. We would like to thank the participants and researchers from the UK Biobank who contributed or collected data. This work was supported by US NIH grants R01DK107859 (to RS), R01HL113338 (to SR), R35HL135818 (to SR), R01DK102696 (to FS and RS), R01DK105072 (to RS and FS), F32DK102323 (to JML), T32HL007567(to JML), K01HL135405 (to BEC), R01HL127564 (to CJW), R35HL135824 (to CJW), and HG003054 (to XZ); Sleep Research Society Foundation Career Development Award 018-JP-18 (to HW); American Thoracic Society Foundation Unrestricted Grant (to BEC); Phyllis and Jerome Lyle Rappaport MGH Research Scholar Award (to RS); UK Medical Research Council MC_UU_00011/3, MC_UU_00011/6 and MR/M005070/1 (to JB, DAL and MNW); UK National Institute of Health Research NF-SI-0611-10196 (to DAL); Diabetes UK grant 17/000570 (to MKR, DAL, and MNW); the Wellcome Trust Investigator Award 107849/Z/15/Z (to DR); Academy of Finland #309643 (to HMO); Stiftelsen Kristian Gerhard Jebsen grant (to KH); Research Council of Norway #231187/F20 (to BW); the Liaison Committee for education, research and innovation in Central Norway grant (to BB); the Joint Research Committee between St. Olavs hospital and the Faculty of Medicine and Health Sciences, NTNU (to KH); the Danish Heart Foundation 16-R107-A6779 (to JBN), and the Lundbeck Foundation R220-2016-1434 (to JBN).

## Author Contributions

The study was designed by HW, JML, SJ, HSD, HO, ARW, VVH, DR, DAW, MKR, MNW, SR and RS. HW, JML, SJ, HSD, HO, ARW, VVH, BB, BW, KK, YS, KP, JB, MAL, FAJLS, SMP, KK, JBN, DAL, MKR, MNW, SR, and RS participated in acquisition, analysis and/or interpretation of data. HW, JML, HSD, HO, KK and RS wrote the manuscript and all co-authors reviewed and edited the manuscript, before approving its submission. RS is the guarantor of this work and, as such, had full access to all the data in the study and takes responsibility for the integrity of the data and the accuracy of the data analysis.

## Data Availability

UK Biobank Sleep Traits GWAS summary statistics are available at the he Sleep Disorder Knowledge Portal (SDKP) website (http://www.sleepdisordergenetics.org).

## Methods

### Population and study design

The discovery analysis was conducted on participants of European ancestry from the UK Biobank study ^27^. The UK Biobank is a prospective study that has enrolled over 500,000 people aged 40-69 living in the United Kingdom. Baseline measures collected between 2006 – 2010, including self-reported heath questionnaire and anthropometric assessments, were used in this analysis. Participants taking any self-reported sleep medication (described elsewhere ^73^) were excluded. 452,071 individuals of European ancestry were studied with available phenotypes and genotyping passing quality control, as described below.

### Excessive daytime sleepiness and covariate measurements

Self-reported excessive daytime sleepiness (EDS) was ascertained in the UK Biobank using the question “How likely are you to dose off or fall asleep during the daytime when you don’t mean to? (e.g. when working, reading or driving)” with the response options of “Never/rarely”, “sometimes”, “often”, “all of the time”, “do not know”, and “prefer not to answer”. Participants reporting “do not know” and “prefer not to answer” were set to missing. Other responses were coded continuously as 1 to 4 corresponding to the severity of EDS. The primary covariates used were self-reported age and sex, and body mass index (BMI) calculated as weight/height^2^. Covariates used in the sensitivity analyses include potential confounders (depression, social economic status, alcohol intake frequency, smoking status, caffeine intake, employment status, marital status, and use of psychiatric medications) and indices of sleep disorders and sleep traits (daytime napping, sleep apnea, narcolepsy, sleep duration, insomnia, and chronotype).

Depression was recorded as a binary variable (yes/no) corresponding to question “Ever depressed for a whole week?”. Social economic status was measured by the Townsend Deprivation Index based on aggregated data from national census output areas in the UK. Alcohol intake frequency was coded as a continuous variable corresponding to “daily or almost daily”, “three or four times a week”, “once or twice a week”, “once to three times a month”, “special occasions only”, and “never” drinking alcohol. Smoking status was categorized as “current’, “past”, or “never” smoked. Caffeine intake was coded continuously corresponding to self-reported cups of tea/coffee per day. Employment status was categorized as “employed”, “retired”, “looking after home and/or family”, “unable to work because of sickness or disability”, “unemployed”, “doing unpaid or voluntary work”, or “full or part-time student”. Day napping was coded continuously (“never/rarely”, “sometimes” or “usually”) responding to the question “Do you have a nap during the day?” Sleep apnea cases (N=5,571) were identified as a union of self-reported and International Classification of Diseases (ICD)-10 coded (G47.3) sleep apnea. Narcolepsy cases (N=7) were determined by the ICD-10 code (G47.4) Insomnia was recorded as “never/rare”, “sometimes” or “usually” responding to the question “Do you have trouble falling asleep at night or do you wake up in the middle of the night?”. Individuals reported “usually” were considered as frequent insomnia symptom cases. Sleep duration was recorded as discrete integers in response to the question “About how many hours sleep do you get in every 24 hours (please include naps)”. In this study, short sleep was defined by sleep duration shorter than 7 hours and long sleep was defined by sleep duration longer than 8 hours. Chronotype was categorized as “definitely a ‘evening’ person”, “more an ‘evening’ than a ‘morning’ person”, “more a ‘morning’ than ‘evening’ person”, and “definitely an ‘morning’ person”. Secondary analyses were performed on participants further excluding shift workers, psychiatric mediation users, and participants with chronic and psychiatric illness (defined elsewhere^73^, N=255,426).

### Activity-monitor derived measures of sleep

Raw accelerometer data (.cwa) were collected using open source Axivity AX3 wrist-worn triaxial accelerometers (https://github.com/digitalinteraction/openmovement) in 103,711 individuals from the UK Biobank for up to 7 days ^80^. We converted .cwa files to .wav files using Omconvert (https://github.com/digitalinteraction/openmovement/tree/master/Software/AX3/omconvert) ^80,81^. Time windows of sleep (SPT-window) and activity levels were extracted for each 24-hour period using the R package GGIR (https://cran.r-project.org/web/packages/GGIR/GGIR.pdf) ^82–84^. Briefly, for each individual, a 5-minute rolling median of the absolute change in z-angle (representing the dorsal-ventral direction when the wrist is in the anatomical position) across a 24-hour period. The 10th percentile of the output was used to construct an individual’s threshold, distinguishing periods with movement from non-movement. Inactivity bouts were defined as inactivity of at least 30 minutes duration. Inactivity bouts with less than 60 minutes gaps were combined to blocks. The SPT-window was defined as the longest inactivity block, with sleep onset as the start of the block and waking time as the end of the block ^85^. We applied exclusion criteria based on accelerometer data quality including 1) none-zero or missing in “data problem indicator” (Field 90002); 2) 0 in “good wear time” (Field 90015); 3) 0 in “good calibration” (Field 90016); 4) 0 in “calibrated on own data” (Field 90017); 5) “data recording errors” (Field 90182) > 788 (Q_3_ +1.5×IQR); and 6) none-zero in “interrupted recording periods” (Filed 90180). Accelerometry data from 85,502 participants of European ancestry passed quality control and were analyzed in this study.

The distributions of accelerometer data are described in Supplementary Table 1. The details of each measurement are as follows. L5 and M10 were the least-active five-hour window and most-active ten-hour window for each day estimated from a moving average of a contiguous five/ten-hour window. The L5 timing was defined as the number of hours elapsed from the previous midnight whereas M10 was defined as the number of hours elapsed from the previous midday. Sleep midpoint was the midpoint between the start and end of the SPT-window. L5, M10, and sleep midpoint variables capture the circadian characteristics of an individual. Sleep episodes within the SPT-window were defined as periods of the z-axis angle change less than 5° for at least 5 minutes ^83^. Sleep duration in a SPT-window was calculated as the sum of all sleep episodes. The mean and standard deviation of sleep duration across all SPT-windows were investigated in this study. Sleep efficiency was calculated as sleep duration divided the total SPT-window duration in a SPT-window. Sleep fragmentation was examined by counting the number of sleep episodes of at least 5 minutes separated by at least 5 seconds of wakefulness within a SPT-window. Diurnal inactivity duration was the total duration of estimated bouts of inactivity that fell outside of the SPT-window in 24 hours, which included both inactivity and naps.

### Genotyping and quality control

DNA samples of 502,631 participants in the UK Biobank were genotyped on two arrays: UK BiLEVE (807,411 markers) and UKB Axiom (825,927 markers). 488,377 samples and 805,426 genotyped markers passed standard QC ^86^ and were available in the full data release. Autosomal SNPs were imputed to a Haplotype Reference Consortium (HRC) panel (∼96 million SNPs). SNPs with minor allele frequency <0.001, genotype calling triplet probability < 0.1, or imputation quality <0.8 were further excluded from the current analyses. The detailed description of genotyping, QC, and imputation are available elsewhere ^86^. We further performed K-means clustering using the principal components (PCs) of ∼100,000 high quality genotyped SNPs (missingness <1.5% and MAF >2.5%) and identified 453,964 participants of European ancestry.

### Genome-wide association analysis

We performed a genome-wide association analysis (GWAS) of self-reported EDS using 452,071 individuals of European ancestry in the UK Biobank across autosomes. A linear mixed regression model was applied adjusting for age, sex, genotyping array, 10 PCs and genetic relatedness matrix, using EDS as a continuous variable of 4 integers, using BOLT-LMM ^28^. Similar linear mixed regression analyses were performed additionally adjusting for BMI and stratified by sex. Secondary GWAS excluding related individuals, shift workers, individuals who used psychiatric medications, and participants with chronic health and psychiatric illness (N=255,426) was performed adjusting for age, sex, genotyping array and 10 PCs in PLINK 1.9 ^87^. Trait heritability was estimated using BOLT-REML ^28^. Genome-wide significance level was set at 5×10^−8^. Gene-sex and gene-health status interaction analyses were performed on unrelated individuals using a linear regression model in PLINK. Conditional analyses to dissect independent signals in significant genomic regions were performed using GCTA-COJO ^88^. Variant annotation was performed using PICS ^58^.

### Post-GWAS analyses

#### Sensitivity and stratification analyses of significant loci

Sensitivity analyses of the genome-wide significant loci in the primary analysis (P<5×10^−8^) were performed additionally adjusting for potential confounders (including depression, socioeconomic status, alcohol intake frequency, smoking status, caffeine intake, employment status, marital status, and psychiatric problems) and clinically important sleep traits (including sleep apnea, narcolepsy, sleep duration hours, insomnia, and chronotype) individually in 337,539 unrelated individuals using PLINK. Sleep traits were further adjusted in the model to investigate their combined effect on EDS signals. Stratified association analyses with EDS were performed in non-obese (BMI < 30, N=256,373) vs obese individuals (BMI => 30, N=81,163), long sleepers (self-reported sleep duration>8 hours; N=25,272) vs short sleepers (self-reported sleep duration<7 hours; N=78,393) and tested for heterogeneity effect.

### Heterogeneity analysis

Genome-wide significant loci identified by primary GWAS analysis were further investigated to understand their contribution to EDS through different mechanisms by testing the associations between EDS risk alleles with BMI, and other sleep traits in the UK Biobank (including self-reported sleep duration, insomnia, chronotype, long sleep duration [>8 hours], short sleep duration [<7 hours], snoring, obstructive sleep apnea defined by ICD-10 code [G47.3], hypersomnolence [defined as sleepiness plus long sleep duration without any chronic or psychiatric diseases], and 7-day accelerometry data). Linear or logistic regression analyses were performed adjusting for age, sex, genotyping array, and 10 PCs. Genome-wide summary statistics of sleep duration, insomnia, chronotype, long sleep duration, short sleep duration, and 7-day accelerometry using BOLT-LMM were available in public database ^73,74,79,89^. We then performed cluster analysis on those loci according to their estimated effect sizes with objectively measured sleep duration, sleep efficiency, sleep fragmentation (number of sleep periods), and insomnia and interpreted their effects as sleep homeostasis or sleep fragmentation.

### Gene, pathway and tissue-enrichment analyses

We further examined the genes within genome-wide significant loci using gene-based, pathway, and tissue enrichment analyses ^63,67,68,90^. Gene-based analysis was performed using PASCAL ^63^. Pathway and ontology enrichment analyses were performed using FUMA ^90^ and EnrichR ^67,68^. Tissue enrichment analysis was performed using MAGMA ^65^ in FUMA, which controlled for gene size. Pathway and tissue enrichment analyses were also performed on genes within loci belonging to sleep propensity and sleep fragmentation clusters separately.

We constructed a weighted GRS comprised of the 42 significant EDS loci and tested for associations with other self-reported sleep traits (sleep duration, long sleep duration, short sleep duration, insomnia, chronotype, and day naps), and 7-day accelerometry traits in the UK Biobank ^73,74,79,89^. Weighted GRS analyses were performed by summing the products or risk allele count multiplied by the effect estimate reported in the primary GWAS of EDS using R package gds (https://cran.r-project.org/web/packages/gds/gds.pdf). We also tested the GRSs of reported loci for insomnia, sleep duration, short sleep, long sleep, day naps, chronotype, restless leg syndrome (RLS), narcolepsy, and coffee consumption associated with EDS using the same approach. The SNPs selected for each trait include 57 genome-wide significant loci for frequent insomnia ^73^, 78, 27 and 8 loci for sleep duration, long sleep, and short sleep respectively ^74^, 348 loci for chronotype ^79^, 125 loci for daytime napping, 20 genome-wide significant loci for RLS ^70^, 6 non-HLA suggestive significant loci (P<10^−4^) in a narcolepsy case-control study of European Americans ^43^, and 8 loci for coffee consumption ^54^.

### Genetic correlation analyses

Genetic correlation analysis using LD Score regression was performed on genome-wide SNPs mapped to the HapMap3 reference panel between EDS (with and without adjustment for BMI) and 233 published GWAS available in LDHub ^72,91^. The significance level was determined as 10^−4^ correcting for multiple comparisons. Pairwise genetic correlations among EDS, frequent insomnia, sleep duration, long sleep duration, short sleep duration and chronotype were performed locally using LDSC. We also partitioned heritability across 8 cell-type regions and 25 functional annotation categories available in LDSC ^92^. Enrichment of the partitioning heritability was calculated in each region with and without extension (±500bp).

### Mendelian randomization analyses

To investigate the causal relationship between EDS and other traits, we performed two-sample Mendelian Randomization (MR) using MRbase package in R ^75^. Inverse Variance Weighted (IVW) approach, assuming no horizontal pleiotropy effect, was implemented as the primary approach in this analysis. BMI, type 2 diabetes, coronary heart disease, psychiatric, reproductive traits, and other sleep and circadian traits (narcolepsy, insomnia, sleep duration and chronotype) were tested as exposures for EDS. Independent genome-wide significant SNPs extracted from publicly available summary statistics of exposures of interest (Supplementary Table 20) were tested as instruments for their effect on EDS. The significance level was determinate as P<0.003 after accounting for multiple comparisons. We identified a putative causal association of higher BMI with increased EDS risk (β=0.018; 95% CI [0.008, 0.028]; P-value=0.0004; Supplementary Table 20). The mean F statistic was 32.7, indicating the instruments were sufficiently strong ^93^. However, Cochran’s Q statistic was calculated to be 677.16 (P=1.09×10^−37^), indicating substantial heterogeneity about the IVW slope. This is an indicator of potential horizontal pleiotropy that violates the traditional IV assumptions. Therefore, we applied MR-Egger regression on the radial plot scale as a sensitivity analysis ^76^. The mean 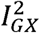 statistic is 0.89, indicating that instruments are sufficiently strong for this analysis ^94^. We observed a consistent effect direction for Radial MR-Egger (β=0.025; 95% CI [−0.005,0.055]; P-value=0.103). Rucker’s Q statistic for Radial MR-Egger is 676.58, indicating that the IVW and Radial MREgger models fit the data equally well (Supplementary Table 21). We also investigated the suggestive putative causal association of type 2 diabetes with increased EDS risk (IVW mean F=29; β=0.005; 95% CI [0.001, 0.009]; P-value=0.014; Supplementary 20). Given variants heterogeneity evidence (Cochran’s Q=88.38; P=0.005), we performed sensitivity analysis using Radial MR-Egger again (mean 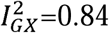) and observed consistent effect direction (β=0.025; 95% CI [−0.005,0.055]; P-value=0.637; Supplementary Table 21). Reverse MR between EDS and other outcome were conducted using 42 genome-wide significant EDS SNPs as instruments, and did not identify any causal association (P>0.05; Supplementary Table 22).

### Replication analyses

#### HUNT

The Nord-Trøndelag Health Study (HUNT) is a large longitudinal population health study, investigating the county of Nord-Trøndelag, Norway since 1984 ^46^. Three surveys (HUNT1 [1984-1986], HUNT2 [1995-1997] and HUNT3 [2006-2008]) have been completed including more than 120,000 individuals. EDS phenotype was collected in HUNT3 by asking the question “How often in the last 3 months have you felt sleepy during the day?” with the choices “Never/seldom”, “Sometimes”, and “Several times”. Individuals with self-reported stroke, myocardial infarction, angina pectoris, diabetes mellitus, hypo- and hyperthyroidism, fibromyalgia and arthritis were excluded from the replication analysis.

DNA samples were collected in 71,860 HUNT samples and genotyped on one of three Illumina arrays: HumanCoreExome12 v1.0, HumanCoreExome12 v1.1 and UM HUNT Biobank v1.0. Imputation was performed on samples with European ancestries using a combined reference panel comprised of the HRC and 2,202 whole-genome sequenced HUNT participants. 29,906 individuals with both phenotype and imputed genotype data were available for this analysis. Sample distributions are presented in Supplementary Table 9a. A generalized linear mixed model analysis was performed on continuous sleepiness adjusted for age, sex, genotyping batch effect, and 4 PCs using SAIGE v0.25 ^95^. A second analysis additionally adjusting for BMI was conducted for replications of loci identified after adjusting for BMI.

#### FINRISK

The FINRISK is a population based study initiated in 1972 and collected every five years since then in Finland to investigate the risk factors for cardiovascular outcomes ^47^. Nine cross-sectional surveys including 101,451 participants aged 25-74 years old were conducted between 1972 to 2012. DNA samples have been collected since the 1992 survey.

We studied exhaustion and fatigue in this population. This was ascertained by asking a question “During the past 30 days, have you felt yourself exhausted or overstrained?” with choises “Never”, “Sometimes” and “Often”. 20,344 individuals with both phenotype and whole genome genotyped and imputed data were available for this study. Genotyping was performed at the Wellcome Trust Sanger Institute (Cambridge, UK), at the Broad Institute of Harvard and MIT (MA, USA), and at the Institute for Molecular Medicine Finland (FIMM) Genotyping Unit using Illumina beadchips (Human610-Quad, HumanOmniExpress, HumanCoreExome). The data was imputed using the 1000 Genomes project phase 3 haplotypes and a custom haplotype set of 2000 whole genome sequenced Finnish individuals as reference panels. Linear regression analyses for exhaustion was performed with snptest v2.5 and adjusted with age, sex, genotyping batch effects and 10 PCs. Shiftworkers were removed from analyses, A secondary analysis was additionally adjusted for BMI.

## Health 2000 Survey

Health 2000 Survey is a population-based sample representing the population structure of individuals from Finland that at the time of contact were over 18 years old. Individuals over 30 years of age answered a number of health and lifestyle-related questionnaires^96^. These data were collected between September 11^th^ in 2000 and March 2^nd^ in 2001 with a goal to reveal and study public health problems in Finland. The Ethics Committee of the Helsinki and Uusimaa Hospital District approved the study protocol, and a written informed consent was obtained from all participants after providing a description of the study. Full Epworth sleepiness scale was included among the questionnaires and included from 4546 individuals with genotyping data on the study.

Genotyping was performed at Finnish Genome center using IlluminaHuman610K genotyping array. Imputation was performed against 2,690 hcWGS and 5,092 WES Finnish genomes (http://www.sisuproject.fi/). Linear regression analyses were performed on continuous ESS adjusted for age, sex, genotyping batch effect and 10 PCs using snptest v2.5. Shiftworkers were excluded and secondary analysis was adjusted with BMI.

A GRS of all 42 EDS loci were also tested in the three replication cohorts. Meta-analysis was performed using Fisher’s method since the measurement of sleepiness were different among the three cohorts.

## Supplementary Materials

Supplementary Tables 1-22

Supplementary Figures 1-9

